# Absolute Quantification of Microbiota in Shotgun Sequencing Using Host Cells or Spike-Ins

**DOI:** 10.1101/2023.08.23.554046

**Authors:** Andrew Wallace, Hong Ling, Sean Gatenby, Seren Pruden, Catherine Neeley, Chad Harland, Christine Couldrey

## Abstract

**Background:** An ongoing challenge for DNA sequencing of samples containing microorganisms is the ability to meaningfully compare different samples and to connect the results back to clinically relevant disease states. The reads of DNA sequence from each sample do not, in and of themselves, give sufficient information to calculate the absolute abundances of each observed organism. Using relative abundances alone is insufficient to determine whether absolute abundances have increased or decreased in the organisms of interest from one sample to the next. This is a well-studied problem in 16S sequencing, but solutions in shotgun sequencing are lacking. Here we show how spike-ins can be used in shotgun sequencing to calculate absolute abundances of organisms present. We also propose the use of the host cells already in the sample as an alternative calculation method. Mammalian host cells are typically of sufficient size that they can be easily and cheaply counted prior to sequencing by a variety of methods and combining this with sequencing data provides sufficient information to calculate the absolute abundances of microbial organisms.

**Results:** Microbial abundances in the samples calculated via this method were consistent with manufacturer-stated values of microbial communities, with qPCR, and with our method tested against itself with regard the spike-in and host-cell based options. *R*^2^ values on the log_10_ scale in these tests ranged from 0.85 to 0.98, and the log_10_-RMSE ranged from 0.1 to 0.7.

**Conclusions:** The proposed method can consistently calculate absolute microbial abundances to within an order of magnitude. Both versions of the method, where spike-ins are added to the samples, or where host cells in the sample are counted, are viable. Calculating absolute abundances allows for direct comparisons to be made between different samples. If disease-thresholds have been identified, absolute abundances can quantify disease states.

## INTRODUCTION

In traditional clinical microbiological settings, the bacterial load is defined in terms of bacterial content per volume of the clinical specimen, e.g. colony forming units (CFU)/ml [1]. However, when performing DNA sequencing of the microbiome it is non-trivial to convert sequencing reads back to this clinically-relevant unit of bacterial load. Data produced by high throughput sequencing runs, be it shotgun sequencing, targeted sequencing, or 16S sequencing, is relative rather than absolute. It is typically presented after taxonomic identification in a format where each genus or species constitutes a given fraction of the reads, with all reads thus summing to 100% for any given sample [2]. It is then easy to compare the relative abundances of microorganisms within a given sample, but meaningfully comparing two such samples is problematic as differences in relative abundances might be caused equally by an increase in the absolute abundance of a genus of interest, a decrease in absolute abundance of other genera, or a combination of factors. Furthermore, these relative abundances do not contain sufficient information to be converted back to the clinically relevant bacterial load.

A substantial number of papers have addressed this issue for 16S sequencing. The read data can be converted to absolute abundances at the genus level if additional measurements are performed on the samples prior to sequencing, such as flow cytometry [2–6], qPCR [7–9], or dPCR [10, 11]. This count then gives a known value for either the total cells or cells of a genus of interest. That information, plus the 16S sequencing output, plus estimations of the number of 16S regions per cell for different genera, allows for an estimate of the absolute abundances of each organism identified in the sequencing data. An alternative method is the addition of a “spike-in” prior to sequencing, as by adding a known quantity of chosen DNA the relative read data can be resolved back to absolute abundances in a similar manner [12–16]. A comprehensive summary of these methods and references is provided by Wang *et al*. [17].

However, no progress appears to have been made towards calculating absolute abundances in shotgun sequencing. In this work we will demonstrate how to adapt the spike-in approach for shotgun sequencing. We also propose an alternative possible approach for calculating absolute microbial abundances based on our work with cow milk but applicable to any samples (e.g. human) that have both host cells and microbes within the sample and where the host cells can be quantified. By counting the number of host cells within the sample via readily available methods (such as somatic cell counting for dairy cow milk), prior to sequencing, the ratio of DNA reads from microbe and host cells then can be used to calculate the absolute abundance of the microbiota of each type using the formulae we provide here.

## MATERIALS AND METHODS

### Sampling

We obtained 393 bulk milk samples from 276 commercial New Zealand dairy herds that conduct periodic milk recording (known as Herd Testing in the southern hemisphere) with Livestock Improvement Corporation (LIC) and which enrolled in our ongoing study of New Zealand dairy milk microbiota. After the animals had been milked into the combined bulk milk vat, farmers collected a sample into a 35 mL vial which contained 0.1mL of 10% concentrate bronopol solution to prevent microbial replication.

On one farm, milk samples from individual cows were obtained on several occasions for DNA sequencing (n=619). These were subsampled from herd test samples that were undergoing routine milk recording at LIC. These samples were heated to 30°C – 36°C and then shaken at 240 rpm for 5 minutes to mix the fat into the sample. An 800 μL aliquot was then subsampled into a 96-well plate for freezing and subsequent DNA extraction and sequencing.

### Laboratory Analysis and DNA extraction

The host cell counts of all samples were measured using standard commercial instruments in the dairy industry (*Bentley FTS Combi* or *Foss MilkoScan FT 6000*). For samples from individual cows this occurred as part of the routine commercial process. For bulk milk samples a 15mL subsample of milk was sent to the Herd Testing facility for host cell counting at the end of each week. All samples were frozen at -20°C until DNA extraction.

Batches of 93 bulk milk samples were thawed until they reached room temperature. Sample vials were then shaken by hand, mixing fat into the milk before aliquoting 1.7 mL whole milk into a 96-well plate and spiking with 17 uL of 1:100 diluted spike-in (*ZymoBIOMICS*^*TM*^ *Spike-in Control I (High Microbial Load)*).

The individual cow milk sample plates were allowed to equalise to room temperature to soften the fat layer before beginning the extraction process. The fat layer was then mixed in by pipetting the samples up and down 3 times. 400μl of whole milk was taken from the herd test subsamples. Samples were then extracted (using a custom *Qiagen BioSprint® 96 DNA Kit* and a *Kingfisher* machine), and eluted with 100 μL of elution buffer. One negative control sample (nuclease free water), and two internal reference *Staphylococcus aureus* qPCR-positive control milk samples were added per plate.

Three further samples were prepared in a similar way, using 325 μL of cow milk taken from three low-SCC animals, adding 75 μL preparations of a mixture of bacteria in known quantities (*ZymoBIOMICS*^*TM*^ *Microbial Community Standard II (Log Distribution)*), and extracting as above. This microbial standard contains 8 bacterial and 2 fungal strains in known quantities in a log-distributed abundance, for use as positive controls in testing microbiome workflows.

### Staphylococcus aureus PCR test

Individual cow milk samples were tested for *Staphylococcus aureus* presence using a commercially available qPCR test from LIC, based on the method of Graber *et al*. [18]. Two *Staphylococcus aureus* PCR positive controls and two nuclease-free water PCR negative controls were used on each 96-well qPCR reaction plate. A dilution series was performed on a positive sample, and CFU counts on blood agar were recorded against CT values from the qPCR to calibrate a CT to CFU mapping.

### DNA sequencing

Short-read (150bp paired) shotgun sequencing libraries were generated using the *Illumina DNA Flex* library prep kit and metagenomic shotgun sequencing was performed on an *Illumina NovaSeq* using *S1* and *S4* flowcells, targeting 15 million reads per sample. Samples were quality checked after sequencing with the *FastQC* program [19]. Samples with fewer than 100k reads were discarded.

### Taxonomic classification

For taxonomic classification, reads shorter than 130bp were discarded, as were reads consisting of all Ns or all Gs (G being the null read in the Illumina chemistry). Reads were identified using *Kraken2* [20], against a database built from microbiome, human, and bovine sequences downloaded from NCBI’s *RefSeq* database [21]. The *Kraken2* ‘confidence’ setting parameter of 0.5 was used to limit false positives in identification.

### Absolute quantification of microbiota

When a whole-cell spike-in is used, estimating the prevalence of a given microbial species (S) of interest is a matter of using the fact that the DNA present from a species in a sample is equal to the number of cells from that species multiplied by its genome length and ploidy. i.e.

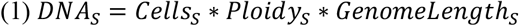

Applying this formula to both the species of interest and the spike-in, and dividing one by the other gives:

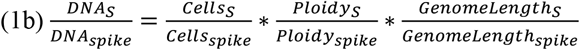

If we ignore, for the moment, any biases that exist in extraction and sequencing, then the sequencer should provide us with reads from both species in proportion to their amount of DNA in the sequencing library. Thus:

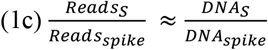

Combining 1b and 1c together and rearranging yields a formula that can be used to calculate the unknown value *Cells*_*S*_from the other known values:

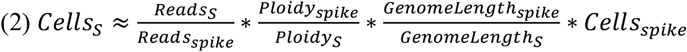

Thus, the known concentration of spike-in cells, can be used in conjunction with the genome lengths and the sequencing reads obtained in the sample to estimate the concentration of cells of a species of interest, S, in the sample. *Cells*_*s*_ and *Cells*_*spike*_ can either be the absolute cell count, or a concentration (e.g. cells/ml), as convenient. We will use units of cells/ml throughout this paper. This formula applies identically to situations where host cells are being used as an internal standard in the sample, rather than a spike-in being added, with the values for the host cells taking the place of the values for the spike-in in the above formulae. In the case where the species of interest is a virus, *Cells*_*s*_ would represent the number of copies of the viral genome present.

In our experience the largest source of error in the above formulae comes from the fact that the *Kraken2* classifier does not resolve all shotgun reads to the species level, but rather assigns reads to the location in the taxonomic hierarchy they best match (Domain, family, genera, subspecies, strain etc) given the confidence setting and sequence database contents. The presence of many similar organisms in the sequence database will reduce how much of a given species’ genome is unique to that species and increase how much is common to other organisms in the sequence database, thus altering the taxonomic classifications of reads from that organism. Thus, the fraction of reads on any given species of interest that are identifiable as having come from that species, varies both by species and *Kraken2* database build. The makers of *Kraken2* provide a program, *Bracken* [22], which can be subsequently run on the *Kraken2* output to approximate a correction for this [23]. Our experience with *Bracken* is that it deals poorly with situations where false negatives are obtained at the species level if the samples have not been sequenced sufficiently deeply to observe nearly all species present, therefore, rather than use *Bracken* for this step in our pipeline, we used the following method of simulating reads on genomes of interest.

To quantify the proportion of reads for each species of interest that were being classified down to the species level we simulated reads on their genomes. We downloaded genome assemblies from *RefSeq* for *Staphylococcus aureus* and *Streptococcus uberis*, used the manufacturer-provided genome sequences for the spike-in bacteria, and used the *ARS-UCD1*.*2* cow genome assembly [24]. We used *RTG-Tools 3*.*10*.*1* [25] to generate 100 thousand simulated reads on each microbial genome and 100 million simulated reads on the cow genome. *Kraken2* was then used to classify the simulated reads, and the proportion of reads classified at the species level or below for each organism in that particular *Kraken2* database was recorded. For the species where the proportion of reads classified at the species level was less than 10%, the proportion classified at higher taxonomic levels were also recorded. (See supplementary material)

We can incorporate this idea into our equations by considering the definition of the proportion of reads classifiable down to the species level for a given species and *Kraken2* database, as:

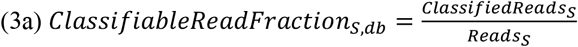

This definition can then be combined with (2) to give:

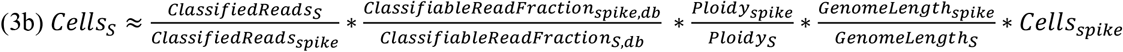

Note that in the case where a portion of the species’ genome is assembled, the relevant genome length to use is the length of the assembled genome and not the estimated length of the total genome of the organism, as reads on unassembled parts of the organism’s genome will neither be able to be simulated nor classified. Also, if the volume of the sample is changed by a spike-in being added to the samples, this formula will give the concentration of species in the sample at the time it was sequenced (i.e. post-spike-in), so additional math will be required to work back to the concentration of species in the sample pre-spike-in.

### Statistical Analysis

The spike in product used (*ZymoBIOMICS™ Spike-in Control I (High Microbial Load)*) contains two bacterial strains, *Imtechella halotolerans* and *Allobacillus halotolerans*. For each of these two spike-in bacteria we used equation 3b to estimate the amount of each the species of interest and took the geometric mean of the results of those two calculations (see Discussion) to obtain an estimated amount of each species of interest in the samples. To calculate the estimated amounts of each species of interest based on the host cell counts, formula 3b was used with host cell values replacing spike-in values in the formula. For estimating total bacterial cells present, an average bacterial genome length of 5 million bp was assumed [26], and it was assumed all reads from bacterial genomes would be classifiable to at least the Domain level. All bacterial abundances are analysed and presented here on a log_10_-scale, as bacteria grow exponentially, and we are dealing with 5-8 orders of magnitude of abundance variation. *R*^2^ values of log-transformed data, and log_10_-RMSE values were calculated in Excel (see supplementary material).

## RESULTS

On average, 19.5 million sequencing reads per sample (SD = 7.6 million) were obtained for the 393 bulk milk samples, with an average of 83% (SD = 1.3) of reads being classified as *Bos taurus* (cow). Seven samples were discarded due to sample degradation, and one sample with <100k reads was omitted from further analysis due to too few reads being obtained. For the 619 individual cow milk samples, an average of 17.4 million sequencing reads per sample (SD = 9.9 million) were obtained.

### Analysis of microbial community standard

*Figure 1* shows the results for the three milk samples spiked with the *ZymoBIOMICS*^*TM*^ *Microbial Community Standard II (Log Distribution)*. After shotgun sequencing of these samples, the amounts of each of the 10 microbes in the community standard was calculated using formula 3b. The calculated amounts of microbes track closely to the manufacturer-stated amounts, with a RMSE on the log_10_-transformed data of 0.55. For the species *Bacillus subtilis*, we report reads based on reads classified at the Family taxonomic level (*Bacillaceae*) as our simulated reads on this species found our *Kraken2* database was unable to resolve reads on this species down to the species level, perhaps due to very closely related species being in the database. These samples were also sequenced without the added product, to confirm that the milk samples themselves did not contain any significant amounts of the same microbes.

**Figure 1:**
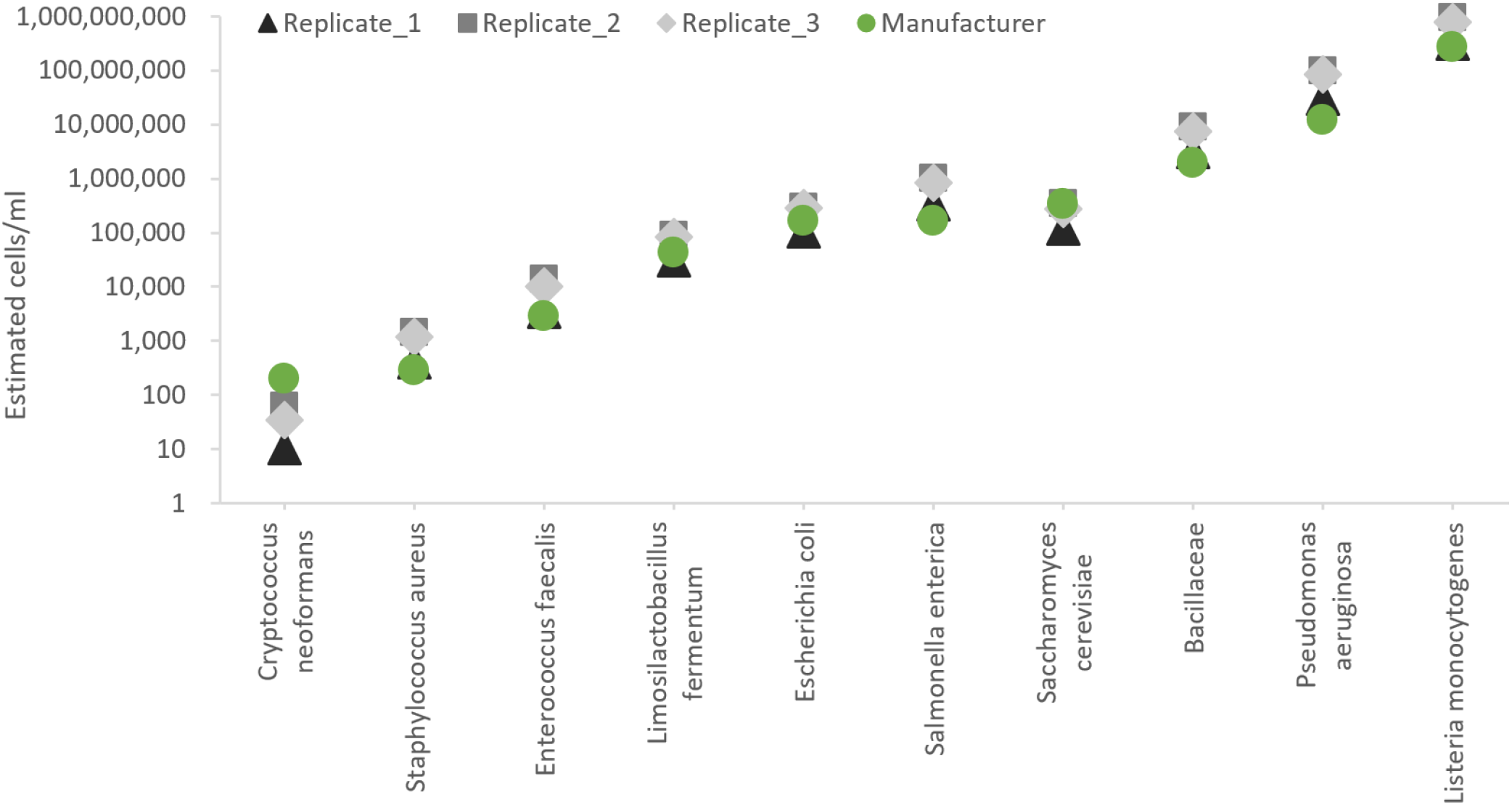
Comparison between manufacturer stated quantities and calculated microbial abundances from shotgun sequencing data. The *ZymoBIOMICS Community Standard II (Log Distribution)* product containing 10 microbial species was added to 3 samples of milk whose host cell count had been measured, and formula 3b was used to calculate the absolute abundances of each microbial organism.

### Comparison with qPCR

On the individual cow milk samples, the commercial *Staphylococcus aureus* qPCR test returned a positive result for 95 of 619 samples. *Figure 2* compares the qPCR abundances of *Staphylococcus aureus* in those samples to the amount calculated from the shotgun sequencing of those samples using equation (3b).

**Figure 2:**
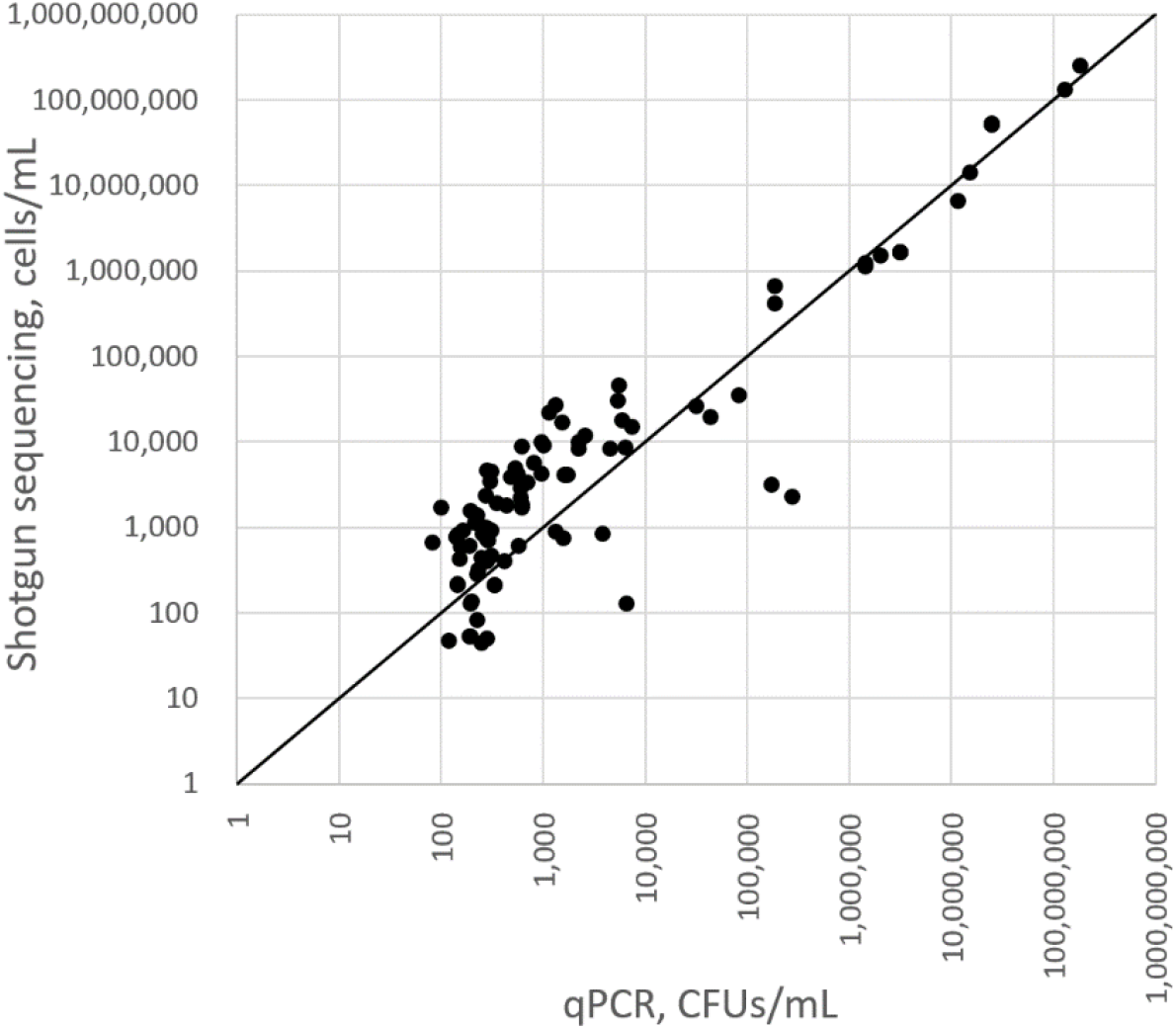
Amount of *Staphylococcus aureus* in individual milk samples, calculated from shotgun sequencing read data using the host cell count of the milk, versus the amount calculated using LIC’s *Staphylococcus aureus* qPCR test. 1:1 line is shown. On the log_10_-scale data R^2^ was 0.85 and RSME was 0.70.

### Comparison of the use of host cell counts with spike-in

The two most common dairy pathogens in New Zealand are *Streptococcus uberis* and *Staphylococcus aureus. Figure 3* shows the amount of these pathogens in the bulk milk samples, and also of total bacteria, as calculated using the host cell count (equation 3b) compared to using a bacterial spike-in. The graphs show strong correlations between the two methods, with *R*^2^ values of 0.91-0.98 at the log_10_-scale.

**Figure 3:**
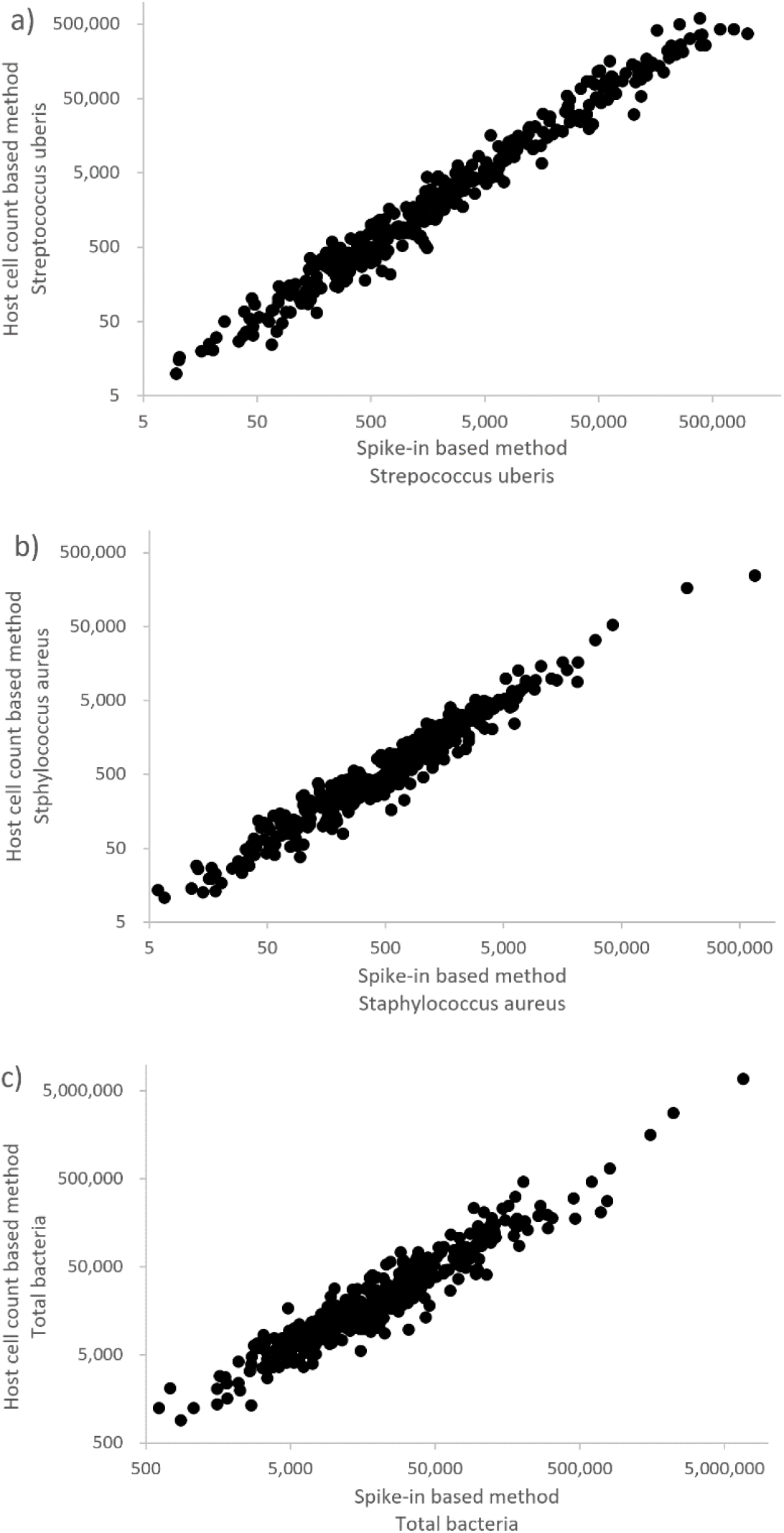
Amount of a) Streptococcus uberis, b) Staphylococcus aureus, c) Total bacteria, in 385 bulk milk samples from New Zealand farms, calculated using the host cell count of the milk, versus the *ZymoBIOMICS* spike-in. *R*^2^ values were 0.98, 0.98, and 0.91 respectively on the log-log scale. Log_10_ RMSE values were 0.13, 0.12, and 0.13 respectively.

## DISCUSSION

The results above show strong agreement of both proposed methods with externally quantified microbial counts, and with each other. The utility of the host-cell count method, as opposed to adding spike-ins to samples will depend on the application. One of the measurements most commonly performed on cow milk in the dairy industry is the measurement of host cell count, known as Somatic Cell Count (SCC). Such measurements are typically performed on individual cows multiple times per year during milk recordings, on bulk milk from the farm multiple times a month by a dairy company, and potentially on the milk from every cow in every milking if the farmer has installed in-line milk-measurement systems. This makes its use for calculating bacterial loads in milk sequencing very attractive, as host cells are already part of the sample, and no external spike-in needs to be purchased and added. Similarly in many clinical settings, human cell counts in some sample types are likely to be readily obtainable. A downside of a host cell count based method is that it cannot be combined with a host-removal method during the extraction process, and so an excessive percentage of sequencing reads may be spent on host cells rather than the microbial cells of interest. When dealing with samples skewed heavily towards host cells, it may be preferable to use bacterial spike-ins for the absolute abundance calculations so that host cell removal methods can be applied during DNA extraction, in order to not spend too many sequencing reads on host DNA. Nor can the host cell based quantification be used in situations where the host cells cannot be counted. For example, in milk and blood the host cells are suspended as single cells and therefore a variety of counting methods can be easily applied, whereas in a tissue sample the host cells may not be able to be quantified.

In our results (see figures 1-2) the ratio between the host-estimated and externally estimated bacterial quantities was always very close to 1. This indicates our extraction method is extracting bacterial and host DNA from cells with close to equal efficiency. In figure 1, *Cryptococcus neoformans* had the largest outliers in relative log values, and we believe this to be due to known difficulties in extracting DNA from *Cryptococcus neoformans* due to it having a thick and resistant capsule accounting for over 70% of cellular volume [27]. The differentially difficult lysability of this species could be accounted for with the addition of an empirically determined multiplier-term *Bias*_*S*_ to the abundance calculation equation 3b. The term would represent a ratio of how easily the host (or spike-in) and the cells of the microbial species of interest are lysed and extracted using the given extraction method, as well as any bias coming from the sequencer (e.g. GC bias). These biases will be constant for a given species, however, and could be ignored for the purpose of comparisons among multiple samples with regard to the relative prevalence of a given species.

It is useful to note that in these equations all terms are multiplicative, so if the interest is always in the same species, most of the terms can be combined in a single unknown *k*. This single *k* value can then be empirically determined from a single sample where the quantities are known, due to a manufacturer-provided quantity, spike-ins, or some other absolute quantification method:

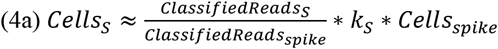

Alternatively the equation can be written in logarithmic form and some use made of the *log(AB/C) = log(A) + log(B) – log(C)* identity to separate some individual terms out. Since bacteria tend to grow exponentially, it will usually make sense to be viewing bacterial counts on a logarithmic scale (e.g. as in figures 1-3). An advantage of a log-form of the equation is that when comparing results for a different species across multiple samples, any unknown terms such as log(*Bias*_*S*_) are simply a constant offset value, so between-sample differences and correlations can be observed even if absolute quantifications are not known precisely as this unknown bias simply has the effect of moving all samples by the same amount along the axis. A disadvantage of separating out too many of the terms into log form, is that *ClassifiedReads*_*S*_ may be zero, and log(0) is undefined. The usual workaround in order to avoid log(0) errors is to add 1 to the term first, but this then introduces some error. In cases where the number of *ClassifiedReads*_*S*_ is generally small across samples, adding 1 to all such terms can skew the result. In order to keep this error to a minimum, we recommend combining as many terms in the equation as possible before finally adding 1 and taking the logarithm of the result rather than separating each and every equation term and only adding 1 to the *ClassifiedReads*_*S*_ value, in order to minimize error introduced in the results.

A question that arises when applying the spike-in methodology to any new sample set is: How much spike-in should be added in the lab? Obviously if too little spike-in is added, it might be that no reads on the spike-in are obtained in some samples and thus no quantification is possible. Whereas if too much spike-in is added to the samples, perhaps >99.9% of shotgun reads obtained from the sample might be on the spike-in, which both wastes much of the money spent on the sequencing, and may reduce ability to see species of interest in the resulting sequence as it may cause some species to drop to zero reads. We suggest the ideal spike-in range to be 100 or above reads obtained per sample on the spike-in, but significantly less than 50% of total sequencing reads per sample. Since samples being sequenced may differ from each other in microbial load significantly, putting the same amount of spike-in into every sample will result in a wide distribution of resulting spike-in read percentages. In our bulk milk samples here, we obtained 7109 reads on average per spike-in species per sample, with the two spike-in species together constituting an average of 0.07% of the reads obtained from each sample. If the obtained spike-in reads for any sample ever drops to zero, then quantification becomes impossible. Similarly obtaining single-digit reads on a spike-in will add a substantial degree of error to the calculations, e.g. obtaining 3 reads by chance on the spike-in rather than 2, will cause a 50% error in calculated cells/ml values for all species.

Due to the exponential reproduction of microbial entities, it is common to see different orders of magnitude of microbes in different samples. In this work bacterial amounts spanned a 10^7^ range of cells/ml. As a result, we have found it always preferable to graph and analyse results on the log_10_ scale rather than the absolute scale. Graphs on the absolute scale of samples spread over orders of magnitude cause most points to bunch around near (0,0) and show only the very highest order of magnitude points. Likewise calculating *R*^2^ and error values on the absolute scale is then equally misleading, as it essentially draws a line of best fit from (0,0) to the highest-magnitude point and reports an almost perfect fit, letting the single high-microbial-load sample dominate all the other samples in its contribution to the calculated error. For this reason, all graphs in this paper have axes on the log_10_ scale, and all reported *R*^2^ values are reported on log_10_-scale data. Deciding what error metric most meaningfully captures error in the data was a non-trivial task. In figures 1 and 3 we can see that the error in the data is generally confined within a certain vertical distance from the truth-set or the 1:1 line on the depicted log-scale graphs. It can be visually observed in these graphs that the amount of error in the y-axis values is similar right across the graph and is not generally a function of the x-axis value. This suggests that RMSE on the log_10_ scale is the best error metric to report. Other error metrics we considered had the significant downside of not being stable across the range of the data, giving higher or lower values for samples of higher vs lower bacterial load. The downside of log_10_-RMSE is that it is not very intuitive in meaning. In figure 1, the log_10_ RMSE was 0.55. This is reflecting the fact that the data generally was about half a y-axis scale-unit of the true value (the y-axis scaling logarithmically in factors of 10). We can calculate 10^log_10__RMSE, ie 10^0.55^ = 3.5, to give a more intuitive meaning for that error amount, as that tells us that the average sample was a factor of 3.5x distant from the truth value. If the truth value was 1000, the samples would thus tend to fall in the range 3.5×1000= 3500 to 1000/3.5 ≈ 286. It is worth noting that this is an asymmetric error range on the non-log scale. It is not truth value ± some error value, and instead the range of error above the truth value is much larger to than the range below the truth value. In the example, the error range is 286 to 3500 for the truth value of 1000, and 1000 is not in the centre of that range. However, on the log scale, it is. The asymmetric nature of the error on the non-log scale as compared to its symmetry on the log scale is worth bearing in mind in any applications of this methodology where accuracy is of great importance (e.g. clinical settings). An important application of this observation is where multiple samples are being taken and then averaged (e.g. to improve accuracy): They should be converted to the log scale and then averaged, as otherwise the asymmetric error will cause the higher samples to dominate. E.g. in this example if two samples were taken and gave 286 and 3500, for the truth value of 1000 as discussed, then the average (arithmetic mean) of those two would be 1,893, almost twice the truth value. Instead putting the values in log_10_, then taking their arithmetic mean, and then raising the result to the power of 10 gives the correct mean of 1000 (this is mathematically equivalent to calculating the geometric 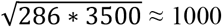. All of this is a result of the formulae given in this paper being multiplicative in nature, and the tendency of multiplicative errors when accumulated to lead to multiplicative error in the result rather than absolute error.

The observation that geometric mean, rather than arithmetic mean, should be used to average multiple samples, was used earlier in calculating the values shown in the figures in this paper. The spike-in used, *ZymoBIOMICS*^*TM*^ *Spike-in Control I (High Microbial Load)*, contains two species, *Imtechella halotolerans* and *Allobacillus halotolerans*. Each of those spike-in species can be used to generate an estimation of the number of cells present from any other species in the sample. Combining those two predictions by geometric mean rather than arithmetic mean consistently gave very slightly better results for the figures in this paper, e.g. a 0.7% reduction in reported log_10_-RMSE error in figure 1; 0.02-0.08% increases in R^2^ values in Figure 3.

Let us consider how the levels of error we observed in our results can arise from errors in the value of the variables. The formulae given are multiplicative, and therefore the final percentage error will be approximately the sum of the percentage error in the individual terms. The percentage error in all individual terms is thus of equal important in the accuracy of the result. We also noted earlier that due to the multiplicative nature of the formulae, in absolute terms the margins of error above the correct value are a lot higher than those below the correct value. Major potential sources of error, apart from the earlier-discussed per-species extraction and sequencing biases, are:

1. The genome length of a given microbiome species of interest. It will commonly be the case that a taxonomic-identification database compiled from publicly available data would include multiple assemblies for the species, including a variety of subspecies, and these assemblies may have different lengths. E.g. one study on *Streptomyces rimosus* found about a +/-10% variation in length compared to the average among the 32 assemblies of the species on NCBI’s *RefSeq*.[28]
2. Randomness in obtained reads resulting from the non-homogeneous nature of the solutions being sequenced. The spike-in product we used contained two bacteria in equal quantities, and we observed a 17% standard deviation in the ratios of these bacteria among our samples, which quantifies the variation due to sample heterogeneity.
3. Inaccuracies in the lab. The sample spiking and extraction process involves a series of pipetting steps with very small volumes, which may have compounding inaccuracies.
4. Inaccurate simulation of the fraction of the reads on a species that fall on the species classification of interest. The program used to generate the reads may not introduce appropriate numbers of errors into the reads that genuinely match the quality of data that would be generated from the sequencer. Or the genomic assembly used as a model from which to generate reads and test species-level-classifiability might deviate substantially from the real-world organism being sequenced.
5. Host cell measurement error. Host cells can be measured in a variety of ways, each subject to different levels of error. The manufacture specifications of our *Bentley FTS Combi* assert a <15% error in our host cell counts.
6. Errors in taxonomic classification. Taxonomic classifying programs can give different results to each other due to differing algorithms and parameters. A significant contributor to variation is also the genetic databases supplied to these programs.

Many of these errors cannot easily be quantified directly, and hence total error cannot be easily predicted and must be observed experimentally.

Despite these various sources of possible error, we saw strong correlations in figures 1 and 2 above when validating the method against qPCR abundances and manufacturer-stated abundances. The method is clearly able to calculate absolute abundances to within an order of magnitude, and this should clearly distinguishes different disease states as we saw the bacterial abundances naturally ranging over seven orders of magnitude. The method provides the ability to directly compare the absolute abundances from different samples for any given species, making data more comparable. The sequencing of all DNA in the sample, and then the calculation of the absolute abundances via this method allows all cellular and viral diseases to be screened for simultaneously, providing a powerful method for health testing.

## CONCLUSIONS

The methodologies outlined here appear to be reliable methods for calculating absolute bacterial amounts from shotgun sequenced samples. This allows for the direct comparison of samples with each other in terms of absolute abundances of the constituent microbial species. This allows the sequenced samples to be meaningfully compared with each other in ways that are not possible with relative abundances alone. It also allows for comparison to known disease thresholds in clinical settings. The use of metagenomic sequencing to screen for all detectable disease genomes simultaneously, and the use of these methodologies to quantify the absolute abundances of the detected disease levels can provide a very powerful health screening tool.

The option of either adding a spike-in to samples, or measuring host cell count, gives a degree of flexibility and both methods are viable. Error appeared to typically be within half an order of magnitude, and that remained true over seven orders of magnitude in the samples analysed here. This allows samples with low microbial loads for a given species to be very clearly distinguished from samples with high microbial loads for that species. In applications where high accuracy is important, repeated samples can be taken and sequenced and the geometric mean used to average them and thus reduce the error, and species-specific extraction bias could be calculated using positive controls in cases where a specific species was of particular interest.

## DECLARATIONS

### Ethics approval and consent to participate

Animal ethics approval was provided by the AgResearch Animal Ethics Committee (AEC), reference 2022-0467-AE-2799. Farmers supplying cow milk samples signed written consent forms.

### Consent for publication

Not applicable.

### Availability of data and material

All data generated or analysed during this study are included in this published article in the supplementary information.

### Authors’ contributions

AW conceived and developed the methodology, performed data analysis, and wrote the manuscript. HL assisted in testing the methodology. SG, SP, and CN performed all lab processing of the samples. CH and CC were involved in initial brainstorming of the methodology and CC provided continuous ideas for testing of the methodology and suggested major revisions to the manuscript. CH and CC wrote the funding application. All authors read and approved the final manuscript.

## Acknowledgements

The authors thank the hundreds of New Zealand farmers who voluntarily contributed samples to this project. The New Zealand eScience Infrastructure (NeSI) facility is thanked for the use of their supercomputing resources. Martina Franz and Grant Anderson are thanked for their feedback on the manuscript and figures. Kathryn Sanders is thanked for helpful discussion and feedback.

